# Evaluating Retention Index Score Assumptions to Refine GC-MS Metabolite Identification

**DOI:** 10.1101/2023.01.26.525730

**Authors:** David J. Degnan, Lisa M. Bramer, Javier E. Flores, Vanessa L. Paurus, Yuri E. Corilo, Chaevien S. Clendinen

**Author notes:** These authors contributed equally (David J. Degnan, Lisa M. Bramer). **Corresponding Author**, *Lisa Bramer.

## Abstract

As metabolomics grows into a high-throughput and high demand research field, current metrics for the identification of small molecules in GC-MS still require manual verification. Though steps have been taken to improve scoring metrics by combining spectral similarity (SS) and retention index (RI), the problem persists. A large body of literature has analyzed and refined SS scores, but few studies have explicitly studied improvements to RI scores. Here, we examined whether uninvestigated assumptions of the RI score are valid and propose ways to improve them. Query RI were matched to library RI with a generous window of +/-35 to avoid unintentional removal of valid compound identifications. Each match was manually verified as a true positive, true negative or unknown. Metabolites with at least 30 true positive identifications were included in downstream analyses, resulting in a total of 87 metabolites from samples of varying complexity and type (e.g., amino acid mixtures, human urine, fungal species, etc.). Our results showed that the RI score assumptions of normality, consistent variance across metabolites, and a mean error centered at 0 are often violated. We demonstrated through a cross-validation analysis that modifying these underlying assumptions according to empirical, metabolite-specific distributions improved the true positive and negative rankings. Further, we statistically determined the minimum number of samples required to estimate distributional parameters for scoring metrics. Overall, this work proposes a robust statistical pipeline to reduce the time bottleneck of metabolite identification by improving RI scores and thus minimizes the effort to complete manual verification.

## INTRODUCTION

Small molecule identification with gas chromatography mass spectrometry (GC-MS) is a powerful approach to understanding the metabolomic composition of samples. Due to its relative inexpensiveness, the usage and thus overall volume of data from GC-MS experiments has increased. In a typical identification workflow, features of GC-MS spectra like retention index (RI) and spectral similarity (SS) are matched to libraries and scored. The current bottleneck in metabolomics experiments is the manual verification of metabolites following identification scoring, as the true metabolite is not always the top rank.^1–6^

Traditionally, SS scores and RI scores have been used to identify metabolites.^3,4,7^ SS calculates the distance between reference and experimental relative mass abundances,^7^ and is a popular approach since spectra provide a vector of datapoints and thus more information for matching metabolites than a single point (like RI). Using SS alone, however, is known to lead to high false discovery rates (FDRs) due to metabolites with similar or the same spectra, called isoforms.^3,4,7^ Isoforms can be distinguished by using retention times (RTs); however, RTs are known to fluctuate across instruments and with experimental parameters like column type, temperature, and age.^8–10^ To address this RT drift, Kovàts et al.^8^ proposed using alkanes to standardize retention times, called retention indices (RIs). Over the last 60 years, RIs have been routinely demonstrated as useful in metabolomic identification,^2–4,6,11–12^ and have been expanded to non-isothermal technologies and other reference compounds, like n-alkanes or Fatty Acyl Methyl Esters (FAMES).^13^ These reference compounds are run and processed with each sample set and facilitate the alignment of elution times within and across studies via RI. The RI score implementation that uses the kernel of the normal distribution has become quite popular.^11,12^ Some popular software, like MetaboliteDetector^11^ assumes the RI distribution is normally distribution for every metabolite and guides users to select a standard deviation (a “RI search window”) between 2-10. This software also assumes the standard deviation for every metabolite within a sample type is known and similar.^11^ MS-Dial^12^ uses the same RI score but does not recommend a standard deviation. Both software combine RI and SS scores to reduce FDRs, but even with these enhancements, the bottleneck of manual verification still persists.^1,2,5^ Any improvements to a single score, SS or RI, will advance scoring in metabolomics overall.

A large body of literature exists regarding the development and advancements of SS scores,^1,3–5,7^ though few studies have focused specifically on developing RI scores,^3,4,11^ especially to datasets outside of the National Institute of Standards and Technology (NIST) Retention Index Library.^13^ Though some have demonstrated that RI distributions are not often normally distributed,^9^ no published study has evaluated the effects of violating RI score assumptions on several complex datasets. Herein, we used both standard and complex samples to evaluate the following assumptions of the RI score: 1) RI are normally distributed, 2) reference retention indices are the means of their prospective RI distribution, and 3) the assumed RI variance is constant across all metabolites. Next, we evaluated approaches to improve the identification efficacy of RI scores. Finally, we used down-sampling and considered the standard errors of distributional parameter estimates to investigate the number of samples, per metabolite, required to characterize a RI distribution. Our approach demonstrates that RI scores which incorporate the properties of each RI distribution outperform those that assume consistent variance, while simultaneously determining a reasonable number of metabolites in which to estimate an RI distribution for a metabolite.

## EXPERIMENTAL SECTION

### Sample Preparation and Data Acquisition

Standard metabolites were purchased from Sigma Aldrich. Dried metabolite standards were derivatized using a modified version of the FiehnLib protocol, as was described in Kind et al.^14^ Briefly, samples undergo methoximation to protect carbonyl groups and reduce tautomeric isomers. This was followed by silylation with N-methyl-N-trimethylsilyltrifluoroacetamide and 1% trimethylchlorosilane (MSTFA) to derivatize hydroxy and amine groups to trimethylsilated (TMS) forms. The samples were then aliquoted into an autosampler tray for GC-MS analysis. An Agilent GC 7890A coupled with a single quadrupole MSD 5975C (Agilent Technologies) was used for collection of GC-MS data. Data was collected over a mass range of 50-550 *m/z*. A standard mixture of fatty acid methyl esters (FAMEs) (C8-C28) was analyzed with samples for RI alignment. The GC oven was held at 60°C for 1 min after injection followed by a temperature increase by 10°C min^-1^ to a maximum of 325°C at which point it will be held for 5 min. All raw data can be found on the MassIVE archive with ID MSV000089933.

### Spectral Matching and Truth Annotation

Query spectra were converted from Agilent .D files to .cdf and matched to our reference database with CoreMS.^15^ Every possible metabolite within RI windows of size −/+35 RI, called an “RI bin”, was returned and subsequently hand-verified to determine true positives. No timewise retention index drift was identified in our samples, which span from 2016 to 2020 (Fig. S1).

In standard mixtures, any small compound that was not included in the mixture was counted as a true negative, except for FAMES and the compounds that can be introduced during sample preparation: carbonate, phosphate, phosphoric acid, glycerol, and propylene glycol. For complex mixtures, other metabolites in RI bins where a true positive was identified were counted as true negatives, except for prevalent sugars, amino acids, nucleic acids, small organic compounds, and the previously stated FAMES and commonly occurring compounds. Additional true negatives were determined using sample type information from the Human Metabolome Database^16^ and CEU Mass Mediator^17^ (fungi) database in a manually curated table (Table S1). All remaining compounds were considered unknown. These datasets with identifications and truth annotations (true positive, true negative, and unknown) can be found at https://doi.org/10.25584/PNNL.data/1902325.

### Data & Software

Two datasets were used for subsequent analyses. The first dataset, hereafter referred to as the “full dataset” was comprised of all RI bins with hand-annotated true positives, their corresponding true negatives, and unknowns for all considered metabolites. A second dataset, hereafter referred to as the “filtered dataset”, was the “full dataset” filtered down to RI bins with metabolites with at least 30 true positive annotations (referred to as the “subset” metabolites). The filtered dataset was restricted to metabolites where enough data was available to characterize the RI distribution and evaluate the underlying assumptions of each retention index distribution. All statistical analysis was done in R version 4.2.1.^18^ All code for plots and analyses are available on Github at github.com/PNNL-m-q/metabolomics_retention_index_score.

### Retention Index Scoring Assumption Assessment

The RI score (1) was calculated for each candidate metabolite, where the *RI_Q_* is the query retention index, *RI_R_* was the reference retention index, and *σ* was a fixed value, typically between 2-10.^11^

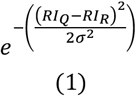

We calculated the effect of varying *σ* on the RI score and rank. As *σ* increases, the median RI score increases for both true positives and true negatives (Fig. S2), but the overall rank does not increase (not pictured). With that in mind, we set *σ* = 3 for all comparisons to the original score throughout this publication.

For each metabolite in the filtered dataset, the Shapiro-Wilks^19^ test was used to assess *RI_Q_* normality, and a Student’s t-test was used to determine if the true average RI error (i.e *μ* = *RI_Q_* – *RI_R_*) was zero. To assess equal variance across *RI_Q_* distributions, an *F*-test was conducted between the metabolites with the lowest and highest estimated variances (*s*^2^). Under the assumption that RI values follow a normal distribution, the mean and standard deviation are thus assumed to be independent of one another, which was investigated with a linear regression model of the RI error standard deviation (*s*) and mean 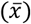.

Then, to understand the effects of violating multiple assumptions on the RI score, we simulated *RI_Q_* – *RI_R_* distributions with every combination of integer mean and skew values ranging from −5 to 5, and standard deviations of 0.5, 1, 2, 3, 4, 5, and 6. These values violated the mean, standard deviation, and normality assumptions simultaneously, as skew was used as a proxy for normality since normal distributions have a skew of 0. If the skew for a particular combination of parameters was 0, the normal distribution sampling function in R was used (rnorm), otherwise, we simulated distributions using a special case of the gamma distribution called the erlang. For target skew values below 0, we multiplied the generated *RI_Q_* – *RI_R_* distributions by negative 1. The average estimation errors for the simulated mean, standard deviation, and skew estimates are 0.0173, 0.0460, and 0.1253, respectively.

### Adjusting Mean and Standard Deviation Assumptions

For each metabolite in the filtered subset, we evaluated the effect of adjusting the assumed *RI_R_* mean and standard deviation with the empirical mean and standard deviation from the observed RI distribution. The performance of RI scores after adjusting for assumptions was evaluated using 10-fold cross-validation. For each fold, 90% of the data was used to estimate the *RI_Q_ s* and 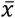 estimates needed to compute adjusted RI scores, and the remaining 10% was used as a validation set to compare assigned ranks (highest score gets rank 1) between the original and adjusted scores. The 10-fold cross-validation procedure was repeated for a total of 50 iterations to account for variability in sampling. Resulting rank changes (new rank subtracted from original rank) were summarized with median and standard deviation rank changes per sample RI bin. Proportions of metabolites with true positives or negatives at ranks 1, 2, and 5 were calculated. To correlate rank improvement with *s*, 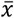, and skew, we binned median rank changes of ≥ 0.05 as “better”, ≤ −0.05 as “worse” and median changes in-between as “no change.” We ran this 10% holdout analysis on the previously described “full” and “filtered” datasets.

### Testing Probability Distribution-Based Scores

After demonstrating the effect of violating the assumptions of the RI score, we tested scores with assumptions that our dataset does not violate. These scores measure the probability a value “fits” within the estimated *RI_Q_* distribution. Here, we simulated the normal, gamma, log normal, and logistic distributions for each subset metabolite. Distributional parameter estimates and their standard errors were obtained using maximum likelihood estimation with the MASS^20^ R package. Theoretical distributions were simulated using the estimated parameters, with 10,000 random draws from the theoretical distribution. To determine the similarity between each estimated and actual distribution, the Kolmogorov-Smirnov (KS) statistic^21^ was calculated between the observed *RI_Q_* distribution and its estimated fit. The best (lowest) KS statistic was identified, and tests for distributional equivalence based on the KS statistic were performed between all estimated distributions. We also computed the Akaike Information Criterion (AIC)^22^ for each estimated distribution, scaling each AIC by subtracting all AICs per metabolite by the lowest value for that metabolite.^23^

New RI scores were calculated by first evaluating the given distribution’s cumulative density function (CDF) at the measured RI. The CDF of a distribution provided the area under the distributional curve at a given point to compute probabilities. If the observed RI was less than the distribution’s mean, the value of the CDF was multiplied by two. If the observed RI was greater than the distribution’s mean, the value of the CDF was subtracted from one and then multiplied by two. Lastly, if it was equal to the mean, a score of one was assigned. This approach measured the probability of observing a retention index that was equal to or more extreme than the observed RI under the assumed distribution. We ran the 10-fold cross-validation analysis 50 times on every score, as described previously, and calculated rank statistics to determine how each score performed in terms of true positives and true negatives. For each fold, the parameters for each respective metabolite’s distribution with the 90% held-in portion of the data were estimated with MASS.^20^ Original RI scores as defined in (1) were used for all metabolites without enough data (less than 30 true positive annotations) to estimate an updated distribution. This analysis was run on both the full and filtered datasets. True positive (TP) and true negative proportions (TN) at rank *n* (1, 2, and 5) were calculated per metabolite as the number of times that molecule was at rank *n* over the total number of respective TP or TN identifications.

### Minimum Requirements to Estimate Retention Index Distributions

After testing scores that use estimations of *RI_Q_* distributions, we then determined a minimum number of samples required to estimate each distribution. For each metabolite in the subset datasets, random samples of sizes 3 to 30 were drawn and fitted to distributional parameters with maximum likelihood estimation.^20^ This sampling process was repeated 100 times for each sample size in the metabolite subset, with all estimates and their standard errors returned. This ensured that all parameter estimates were calculated on the same sampled metabolites per iteration. Distributions of standard errors from each sample size and metabolite scaled to their respective estimates were calculated.

## RESULTS AND DISCUSSION

### Truth Annotation

Across 4,523,251 candidate matches, 13,487 were hand-verified as true positives and 610,403 were determined to be true negatives. The remaining candidate matches were considered “unknown” due to the high presence of potential matches in complex samples where all possible compounds are not fully characterized. In terms of samples, we had 204, 103, 89, 66, 30, and 16 of standards, human cerebrospinal fluid (CSF)^25^, human blood plasma^25^, human urine^25^, fungi species (*A. niger*, *A. nidulans*, *T. reesei*), and soil crust, respectively (Fig. 1a). Across sample types, the mean true positive rate across complex samples was relatively consistent (between 29-37) as opposed to standards with a mean of 4.2039. These rates demonstrated that though it was time-consuming to manually annotate complex samples, these datatypes are advantageous for statistical analyses due to their higher number of true positive annotations on average.

**Figure 1.**
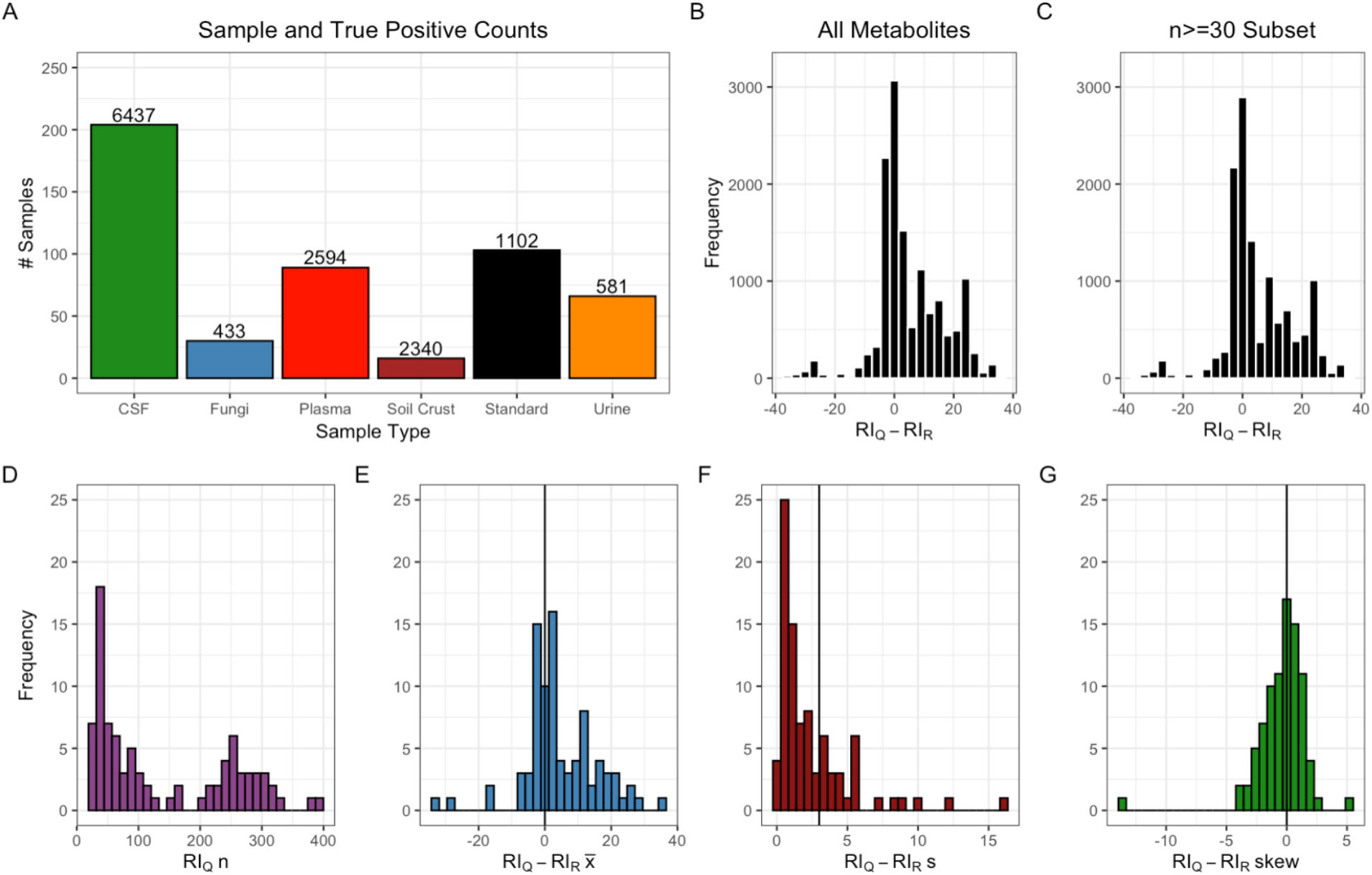
Summary statistics for true positives. (A) Sample counts per type with the number of true positive metabolites written above each bar in text. (B) Distribution of query retention index minus reference retention index (*RI_Q_* – *RI_R_*) for all true positive metabolites and the (C) n ≥ 30 subset. (D) Frequency of metabolite counts in C. (E) For metabolites with a true positive count of at least 30, each *RI_Q_* – *RI_R_* distribution is plotted with a black line to indicate the assumed values of the mean, (F) standard deviation, and (G) skew. A normal distribution assumes as skew of 0.

The subset dataset consisted of a representative subset of 87 metabolites, out of 221 metabolites identified, with at least 30 true positives. The subset dataset was used to calculate summary statistics of the *RI_Q_* distributions (Fig. 1b-c). The number of samples per metabolite in the subset ranged from 30 to 398 with a median count of 88 (Fig. 1d). Mean retention index difference values (*RI_Q_* – *RI_R_*, henceforth referred to as *RI_D_*) spanned the CoreMS^1^ retention index search window of −/+ 35 RI, with a median at 1.86 instead of the assumed value of 0 (Fig. 1e). The null hypothesis that the mean *RI_D_* was equal to 0 for each metabolite was rejected for 85 of the 87 metabolites in the subset using a Type 1 error rate of *α* = 0.05. Though the assumed value was typically in the range of 2-10,^11^ the observed *RI_D_* standard deviation values ranged from 0.01 to 16 with a median value of 1.37 (Fig. 1f). An *F*-Test between the lowest and highest *s*^2^ provided evidence against the null hypothesis of equal variances across metabolite *RI_D_* distributions (*p* < 0.0001). The skew of the *RI_D_* distributions varied from −13.37 to 5.45 with a median of ~0 (Fig. 1g), but none of the *RI_D_* distributions passed the Shapiro-Wilks test for normality under an assumed Type 1 error rate of *α* = 0.05. Given these analyses, the RI score assumptions of normality where the distribution was centered at *RI_R_* and the variance was equal across sample type are likely not true for all metabolites.

### Violating RI Score Assumptions

We simulated the effect of violating one of the mean, standard deviation, and normality assumptions using the summary statistics from our measured *RI_D_* distributions. Skew was included as one example of violating a normality assumption, as the normal distribution was assumed to have a skew of 0. When no assumptions are violated, the median RI score was ~0.8 with an inner quartile range of 0.52 – 0.95 (Fig. 2a). Median RI scores quickly dropped off when violating only the mean assumption, as any |*RI_D_* mean| > 5 returned a median score less than 0.25 (Fig. 2b). This was particularly concerning given that our *RI_D_* 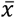 values range from −35 to 35. Interestingly, standard deviation values below 3 tended to have median scores above the no violation distribution, and the opposite was true for standard deviations above 3 (Fig. 2c). In other words, at smaller standard deviations, the RI score was more tightly centered about 0, the true mean. Consequently, a larger majority of the observed *RI_Q_* from this distribution had scores closer to 1. The reverse was true as the standard deviation increases: a smaller majority will be near the true mean and the computed RI scores will be further from 1. Any skew larger or smaller than 0 tended to have higher median scores than the no violation distribution, with |*RI_D_* skew| > 3 resulting in RI score medians near 1 (Fig. 2d). Violating two or more assumptions had powerful combinatorial effects on the retention index score (Fig. 2e). For example, selecting a mean, standard deviation, and skew of 0, 1, and 0, respectively, gave a median RI score 0.9748, while selecting 5, 6, and 0 gives 0.1770, which was more than an 80% reduction in value.

**Figure 2.**
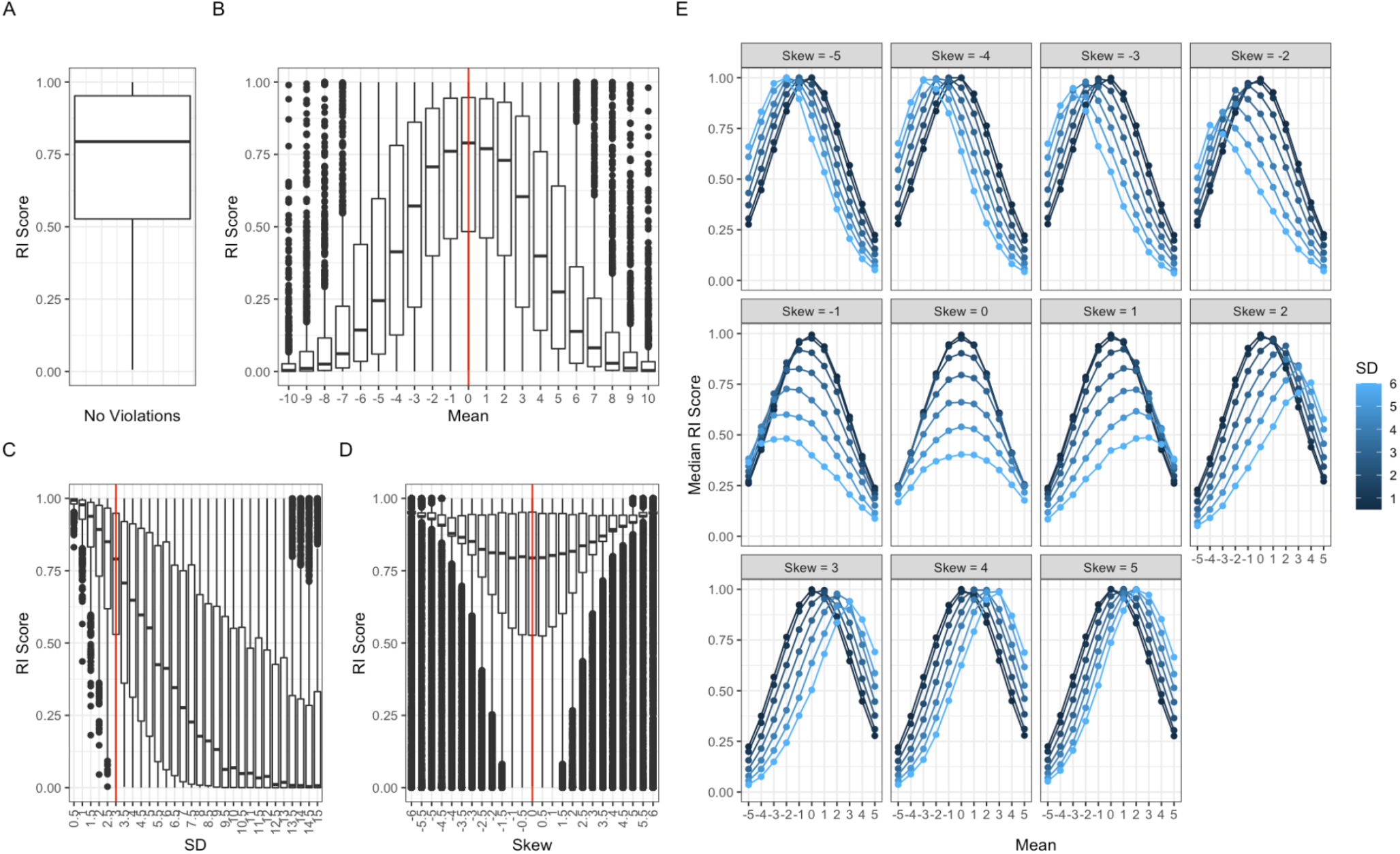
The effect distributional parameters have on the RI score. (A) Distribution of retention index scores when drawing from the assumed retention index query distribution. (B) The effect adjusting only the mean, (C) standard deviation, and (D) skew has on the retention index score distributions, with assumed values marked in red. (E) The effect that adjusting mean, standard deviation, and skew combinatorically has on the median retention index score (n = 10,000 for each simulated distribution, indicated by a point).

### Adjusting the Original RI Score

In the cross-validation analysis on the full dataset, adjusting the RI score to the query 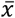 and *s* improved both true positive and true negative average proportions at ranks 1, 2, and 5 (Fig. 3a). Studied ranks had at least a 2-fold improvement in TP average proportions, with rank 1 having the most notable improvement (4-fold, 0.05 to 0.20). TN average proportions were similar or decreased even at ranks as high as 5.

**Figure 3.**
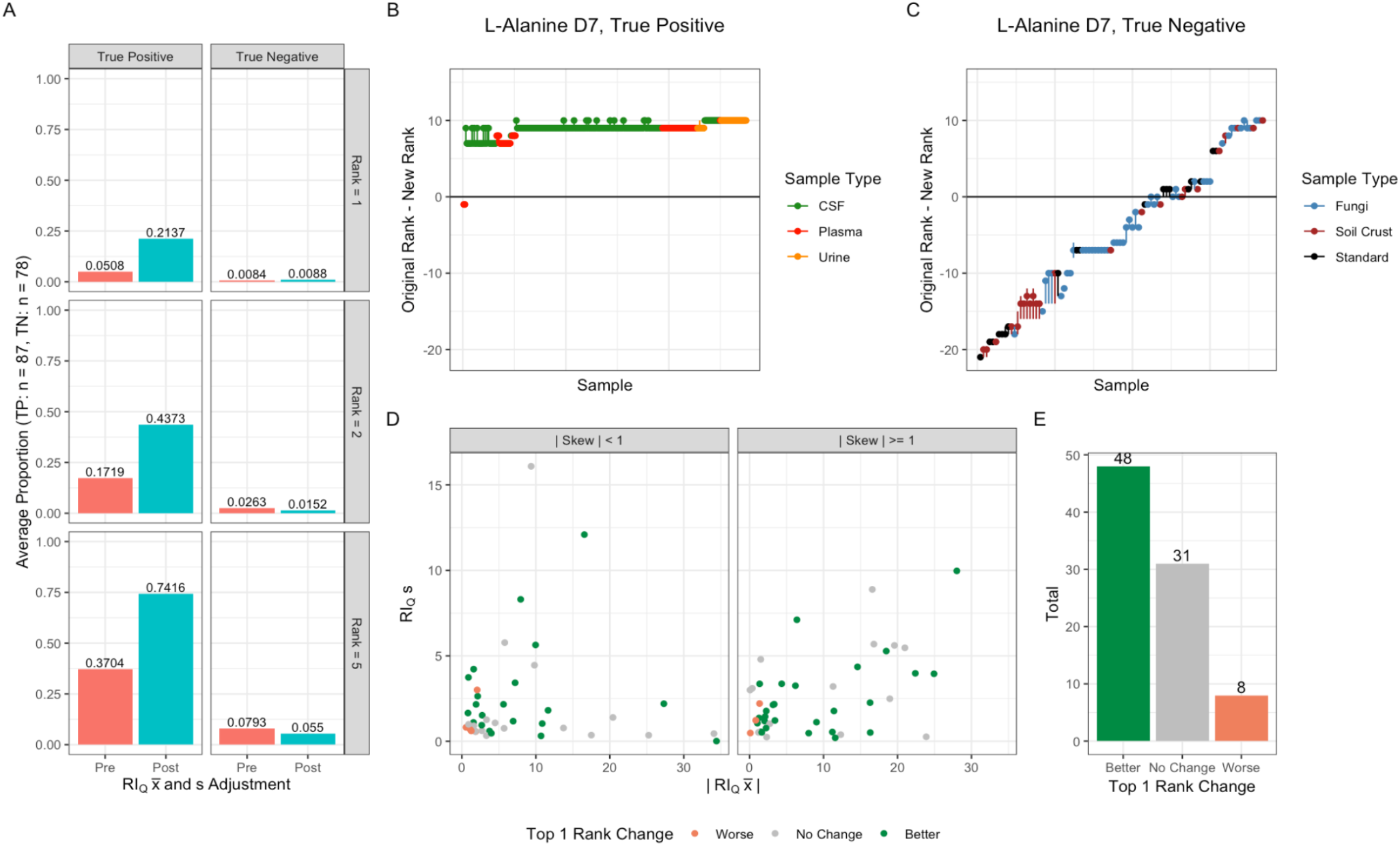
Performances of the adjusted RI score, where the reference retention index was the mean of each metabolite’s distribution, and the retention index search window (standard deviation) was the standard deviation of each metabolite’s distribution. (A) Average proportion of true positives (n = 87) and true negatives (n = 78) at ranks 1, 2, and 5. (B) Distribution of rank changes (new rank subtracted from original rank) for L-Alanine D7 following 10% holdout analysis, colored by sample type and ordered on the x-axis from smallest to largest original rank. Points indicate the median of the distribution, while lines indicate the range for both true positives and (C) true negatives. (D) True positive retention index difference (query minus reference) standard deviations plotted against absolute means, faceted by skews greater than or equal to one. Points are colored by rank change improvements at rank 1, where better ≥ 0.05, worse is ≤ −0.05, and the rest are labeled no change. Counts of these changes are summarized in (E).

To illustrate the results of the cross-validation analysis on the full dataset, we plotted each metabolite’s distribution of rank change (new rankings subtracted from original rank) for true positives (Fig. 3b) and true negatives (Fig. 3c) and have selected L-Alanine D7 as an example. For most metabolites, we saw improvement in the true positives (positive rank change) and true negatives (negative rank change), though the effect that sample type has on rank improvement is not well understood. For example, plotting the rank variation per iteration of the cross-validation analysis per sample type clarified that this variation in ranking was not consistent per sample type, regardless of whether the identification was a true positive or negative (Fig. S3). Though we knew that the mean, standard deviation, and skew influence the RI score (Fig. 2e), the relationship each parameter had on the true positive (Fig. 3d) and true negative (not shown) rank improvements was not apparent for this dataset. Overall, we saw improvement in most metabolite identifications with this approach (Fig. 3e), where we adjust the original score to reflect each *RI_D_* 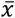 and *s*.

### Parameter Estimation and Fits of Probability Distributions

Using maximum likelihood estimation with the MASS R package,^20^ parameter estimates for the normal, gamma, log normal, and logistic distributions were obtained based on fits to the observed *RI_Q_* distribution. Parameters for all distributions were successfully estimated apart from the logistic parameters for L-proline and arabitol for the full dataset. Several parameters for the gamma distribution failed to estimate a standard error and were indicated in red (Fig. S4).

We calculated Kolmogorov-Smirnov (KS) statistics between the empirical *RI_Q_* distribution and each of its parametric fits. A lower KS statistic is indicative of a better fit,^22^ and the average KS statistic across the 87-metabolite subset was 0.229, 0.264, 0.270, and 0.543 for logistic, log-normal, gamma, and normal, respectively. Tests based on the KS statistic were performed between the best fitting parametric distribution and each other fit to determine their equivalence (Fig. 4a). The total count of best fits for each score was 64, 12, 10, and 1 for logistic, log-normal, gamma, and the original score’s fixed mean and standard deviation values, respectively. The normal distribution was never identified as the best-fitting distribution.

**Figure 4.**
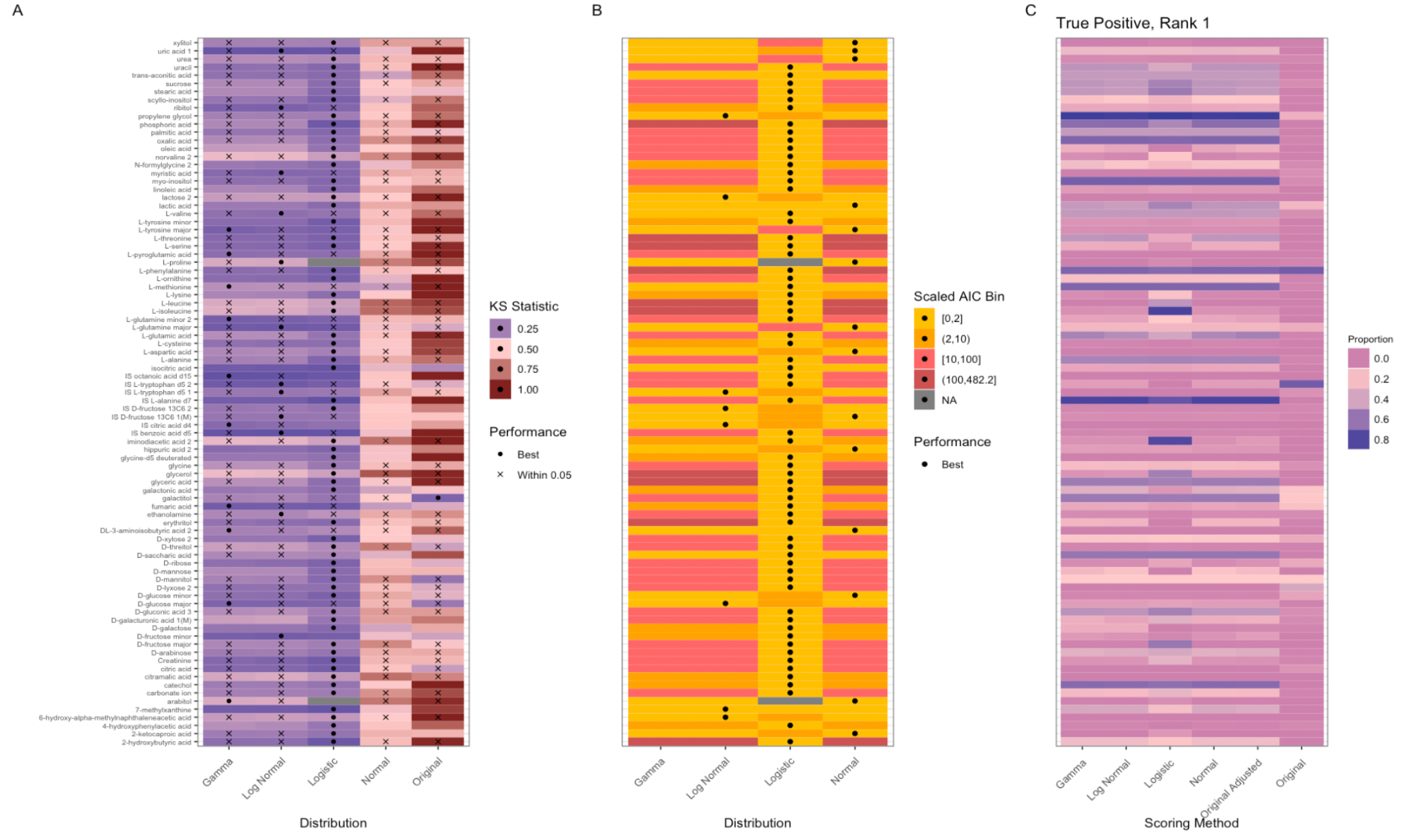
Fits and performances of tested scores. (A) The Kolmogorov-Smirnov (KS) statistic of the distance between the query retention index (*RI_Q_*) distribution and a modeled *RI_Q_* distribution using maximum likelihood estimation. The labels refer to a distribution with the lowest KS statistic (“best”), or one with a p-value smaller than 0.05, which are non-equivalent fits to the best. Across all heatmaps, gray is indicative of a distribution that failed to estimate parameters. (B) The Akaike Information Criterion (AIC) statistic of the distance between the *RI_Q_* distribution and an estimate of that distribution using maximum likelihood estimation. AIC has been scaled to the minimum AIC per metabolite, and then binned. (C) Proportion of samples per metabolite with a true positive at rank 1 for all tested distributions, following a 10% holdout calculated 50 times. “Original” refers to the metabolite detector score where the mean is 0 and the standard deviation is 3, whereas “Original Adjusted” uses the metabolite detector score with each *RI_Q_* estimated mean and standard deviation. All other labeled scores correspond to the probability scores based on the CDFs of the indicated distributions (i.e. Normal, Gamma, Log Normal, and Logistic).

Another metric to indicate the quality of a distributional fit is the Akaike Information Criterion (AIC). To make equitable comparisons, we subtracted each AIC by the minimum AIC per metabolite and counted that minimum and values within 2 of the minimum (i.e. the best fitting in the group) as having equitable fits.^24^ The total count of equitable best fits was 68 for logistic, and 27 for normal, log normal, and gamma. Generally, wherever logistic was not the best fit, the other three distributions were equivalently better fits and vice versa, with some exceptions. Overall, both the KS statistic and AIC have suggested that the logistic distribution was a better fit to the empirical *RI_Q_* distributions (Fig. 4a-b).

### Performance of Probability-Based RI Scores

In terms of true positives, the modified original score (which uses *RI_Q_* mean and standard deviation, instead of the assumed values) and the four probability-based scores all outperformed the original score on average at ranks 1 (Fig. 4c) and 5 (Fig. S5) for both the full (Fig. S4, Table 1) and the filtered (Fig. S6, Table S2) datasets. In terms of true negatives, the new scores were not worse with true negatives than the original at rank 1. The new scores performed better at ranks 5 for the full dataset (Table 1), and performed better at rank 1 and equitably with rank 5 for the filtered dataset (Table S2). The differences in results between the full and filtered datasets can be explained by larger average counts of candidate metabolites per RI bin in the full dataset (56.1 as opposed to 5.71 in the filtered). When the bin sizes were larger, there was more room for improvement in terms of true positives and true negatives simply by the existence of more ranks within an RI bin. We saw that in smaller bins, the proportions of true positives and true negatives were larger than in the full dataset’s larger bins. Ranks as high as 5 often included the entire RI bin in the filtered dataset, which was why true negatives at rank 5 were much higher in the filtered dataset than the full. Overall, the tested scores performed better than the original RI score on average, which highlighted two key ideas: 1) universally assuming every *RI_Q_* distribution was the same was incorrect and modifying for these distributional properties improved score performance, and 2) current reference RI values may need modification by continued experimentation, potentially on a per instrument and column basis.^9,10^

**Figure 5.**
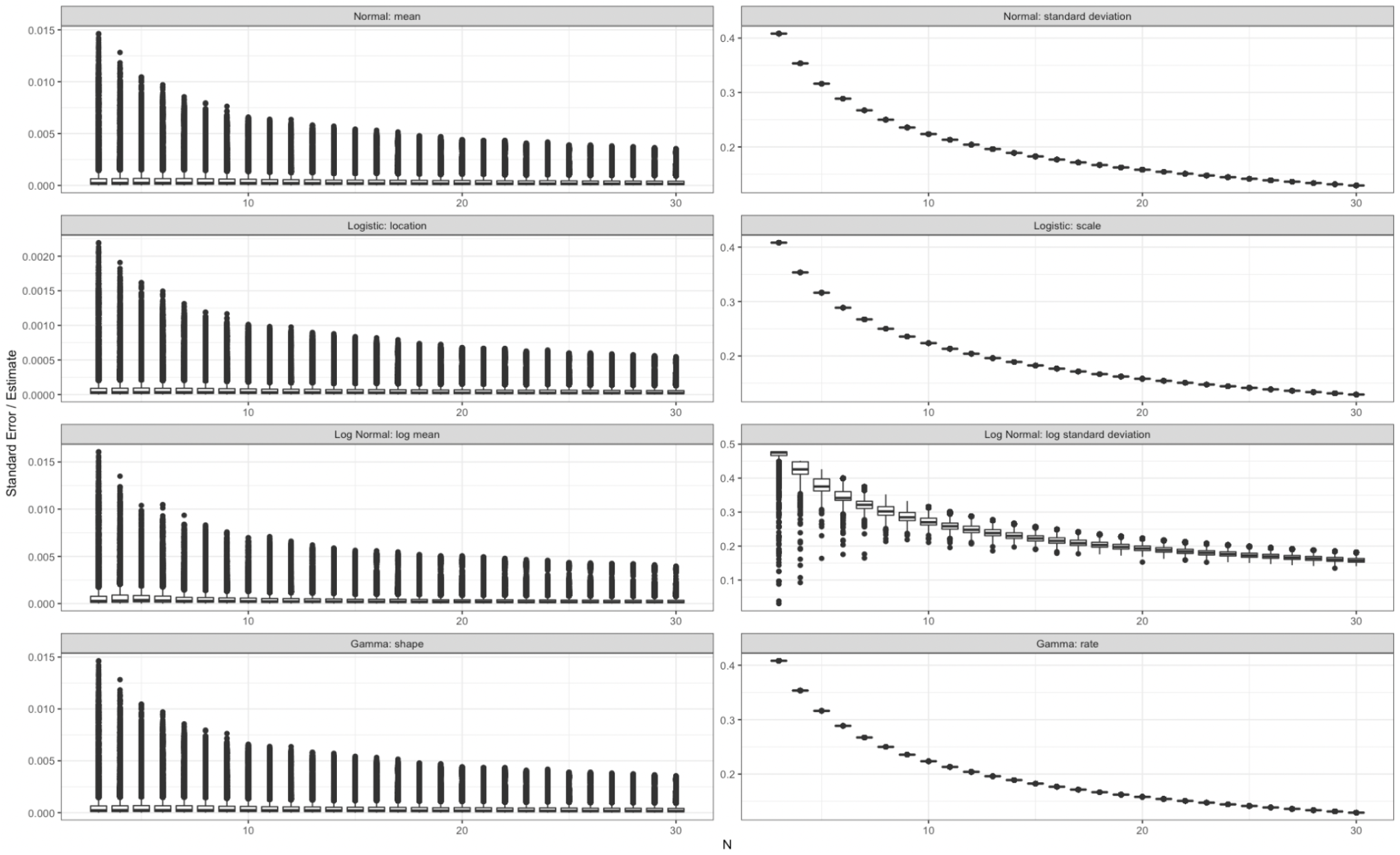
Distribution of the standard error over the parameter estimates following sampling sizes of 3 to 30 repeated 100 times for 87 metabolites, for the normal (top row), logistic (2^nd^ row), log normal (3^rd^ row), and gamma (last row) distributions.

**Table 1.** Average proportion of true positives or true negatives at the specified rank for each score following the 50 iterations of the 10% holdout analysis with the full dataset.

| *Truth and Rank* | *Original* | *Original Adjusted* | *Normal* | *Gamma* | *Log Normal* | *Logistic* |
| --- | --- | --- | --- | --- | --- | --- |
| *True Positive, Rank 1* | 0.0508 | 0.2137 | 0.2132 | 0.2151 | 0.2129 | 0.2418 |
| *True Positive, Rank 5* | 0.3704 | 0.7416 | 0.7414 | 0.7422 | 0.7404 | 0.7002 |
| *True Negative, Rank 1* | 0.0084 | 0.0088 | 0.0087 | 0.0086 | 0.0087 | 0.0092 |
| *True Negative, Rank 5* | 0.0793 | 0.0550 | 0.0546 | 0.0558 | 0.0548 | 0.0630 |

Interestingly, comparing these results to the best fitting distribution according to the KS and AIC, logistic performed the best in terms of true positives at rank 1, but not at rank 5 nor in terms of true negatives (Table 1 and Table S2). This was because the other alternative scores perform relatively equitable to each other, and better than the original score. A downside to using a logistic score was that the maximum likelihood estimation may fail to estimate parameters, as logistic failed to estimate in more than 50% of the iterations of the cross-validation analysis for 4 metabolites within the filtered dataset (Fig. S6).

There were metabolites where the original score outperformed the other metrics in terms of true positives and true negatives at ranks 1 and 5 (Table 2). These cases tended to be at *RI_D_* |*x*| < 2 and *RI_D_ s* < 3 (Fig. S7, full dataset) for true positives, and *RI_D_* |*x*| > 2 and *RI_D_ s* > 3 for true negatives. This was consistent with our observed decreased performance of the original score at means near the assumed value and standard deviations at or below the assumed value (Fig. 2b,c,e). Skew did not appear to play much of a role in whether the original score performed better than the others for a specific metabolite (Fig. S7). This was not surprising, as the original RI score performed better with more extreme skews (Fig. 2d-e). Overall, the original score performed better when the *RI_Q_* 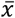 and *s* were closer to their assumed values for true positives, or conversely, further away for true negatives (Fig. S7). This trend was emphasized when we overlay estimated and measured *RI_Q_* distributions for metabolites where the best performer was the original MetaboliteDetector/MSDial score^11,12^ or the other estimated distributions for true positives or negatives at rank 5. The cases where the original score performed the best out of all the scores were where most of the measured *RI_Q_* distribution fell within the assumed standard deviation of the reference retention index value for both true positives and negatives (Fig. S8). Other scores performed better when their estimated distributions overlap more of the measured *RI_Q_* than the original score (Fig. S8).

**Table 2.** Counts of metabolites within our subset (n = 87) where the original or any other tested score has the highest (true positive) or lowest (true negative) proportion of metabolites at ranks 1 and 5. Ties between the original and other score, and cases where all proportions are 0 are also counted.

| *Truth and Rank* | *Original* | *Other* | *Ties* | *All 0* |
| --- | --- | --- | --- | --- |
| *True Positive, Rank 1* | 10 | 58 | 11 | 8 |
| *True Positive, Rank 5* | 18 | 36 | 32 | 1 |
| *True Negative, Rank 1* | 6 | 1 | 18 | 62 |
| *True Negative, Rank 5* | 7 | 9 | 30 | 41 |

### Estimation of RI Query Distributions

After establishing that distribution-based scores perform better on average than the original score, we investigated the minimum number of samples required to estimate an *RI_Q_* distribution. The standard errors (SEs) scaled to their respective parameter estimates tended to decrease as the number of samples increased, as expected (Fig. 5). Across all parameters, the SE followed a curve that tends to stabilize around 10 metabolites, suggesting that a sample size of 10 may be a bare minimum number of runs required for estimating a metabolite’s retention index distribution (Fig. 5). Acquiring more samples than 10 is preferable, as an increase in sample size tended to decrease SE, especially in parameters that model distributional variability (e.g., standard deviation, scale, and rate).

## CONCLUSIONS

Here we demonstrated that there is room for improvement in RI scores and proposed approaches to improve the ranking of metabolites, thus reducing the amount of manual vetting required in metabolite identification. The underlying assumptions of the retention index score regarding the mean, standard deviation, and normality were violated by nearly all metabolites in our dataset, and these violations greatly reduced the RI score of true positives. We tested five alternative scores that attempted to correct for some of these violated assumptions and all performed as well as or better than the original metric in terms of true positive and negative rates. Using the standard errors of estimates, we determined a minimum number of samples to estimate *RI_Q_* to be about 10, with the understanding that more samples will yield better results. Based on this analysis, a potential pipeline for improved RI scores would 1) store the results of all verified metabolite RI, 2) estimate parameters for a preferred alternative score based on a set minimum number of metabolites (10 or greater), and 3) apply the scores and save the verified RI if desired. Overall, this work demonstrated that an RI score that incorporates *RI_Q_* distributional properties will increase the ranks of true positive metabolites, thus generating more accurate small molecule identifications and reducing manual verification efforts.

## ASSOCIATED CONTENT

### Supporting Information

Supporting Information File 1 – Supplemental Figures (PDF). Supporting Information File 2 – Supplemental Tables (XLSX).

## AUTHOR INFORMATION

### Author Contributions

VLP prepared and conducted GC-MS sampling; YEC developed and ran the metabolite identification algorithm; DJD, LMB, JEF developed the statistical analysis pipeline; CSC provided the truth annotations and supervised the project; and DJD wrote the manuscript with significant contributions from all authors. All authors have given approval to the final version of this manuscript.

### Funding Sources

This work was supported by the PNNL Laboratory Directed Research and Development program, and is a contribution of the *m/q* Initiative. The authors declare no competing financial interest.

## ACKNOWLEDGMENT

The authors would like to thank Allison Thompson and Jordan Rabus for their computational assistance in this project.

---

For TOC Only

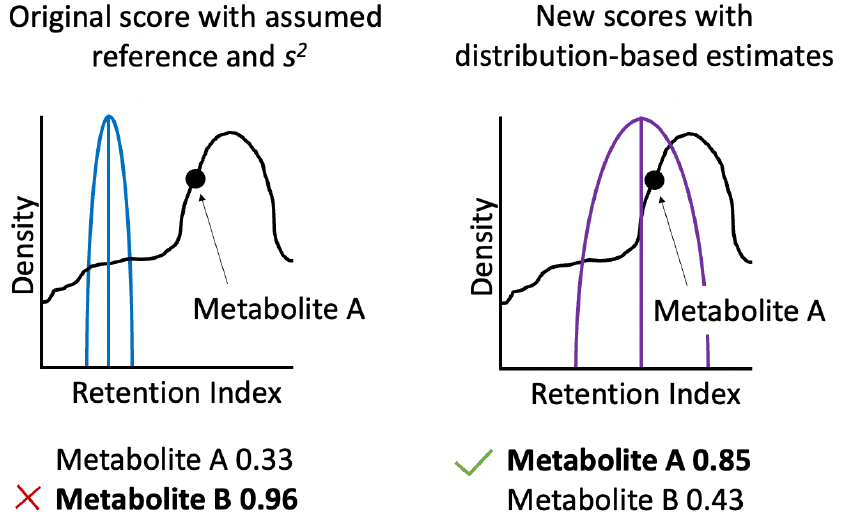

## Notes

https://github.com/PNNL-m-q/metabolomics_retention_index_score

https://data.pnnl.gov/group/nodes/dataset/33302

### Competing Interest Statement

The authors have declared no competing interest.

