## Supplemental Data 1 for "Evaluating Retention Index Score Assumptions to Refine GC-MS Metabolite Identification"

### Table of Contents

| | | |
|---|---|---|
| Supplemental File S1. | S1_Supplemental_Figures.docx | Contains all supplemental figures |
| Supplemental File S2. | S2_Supplemental_Tables.xlsx | Contains all supplemental tables |

### Contents


| **Object** | | **Page** |
|---|---|---|
| Figure S1 | ………………………………………………………………… | S3 |
| Figure S2 | ………………………………………………………………… | S4 |
| Figure S3 | ………………………………………………………………… | S5 |
| Figure S4 | ………………………………………………………………… | S6 |
| Figure S5 | ………………………………………………………………… | S7 |
| Figure S6 | ………………………………………………………………… | S8 |
| Figure S7 | ………………………………………………………………… | S9 |
| Figure S8 | ………………………………………………………………… | S10 |

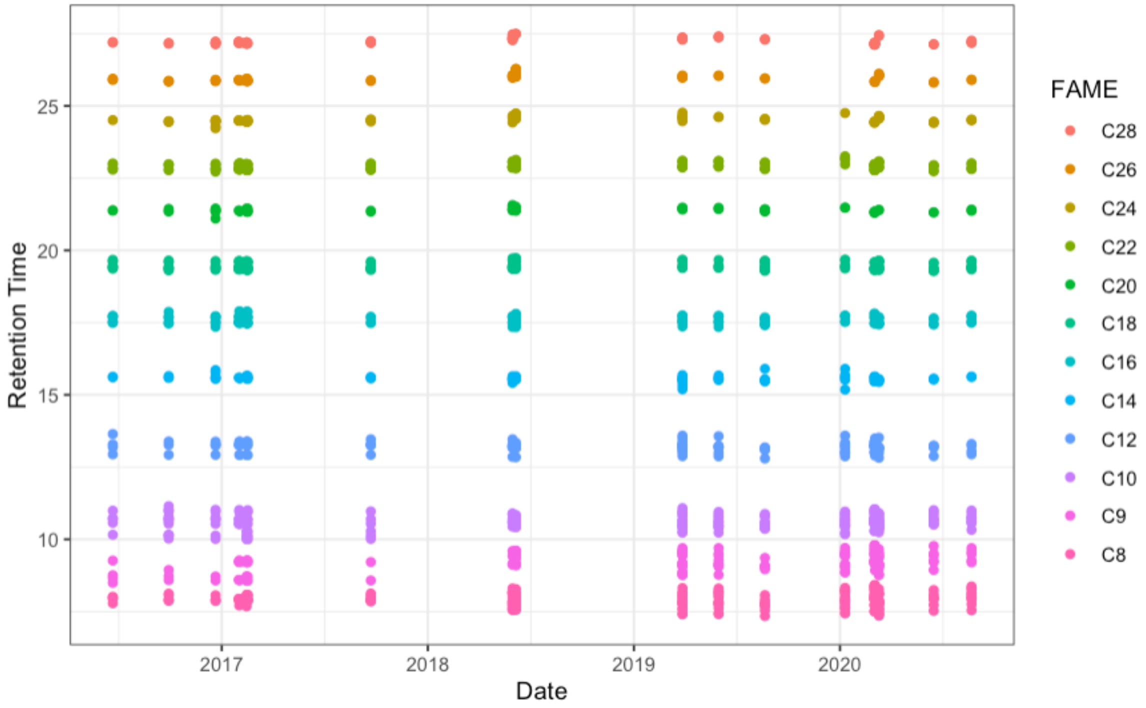


**Figure S1.** Retention times for identified fatty acyl methyl esters (FAMEs) across all samples (n = 548).

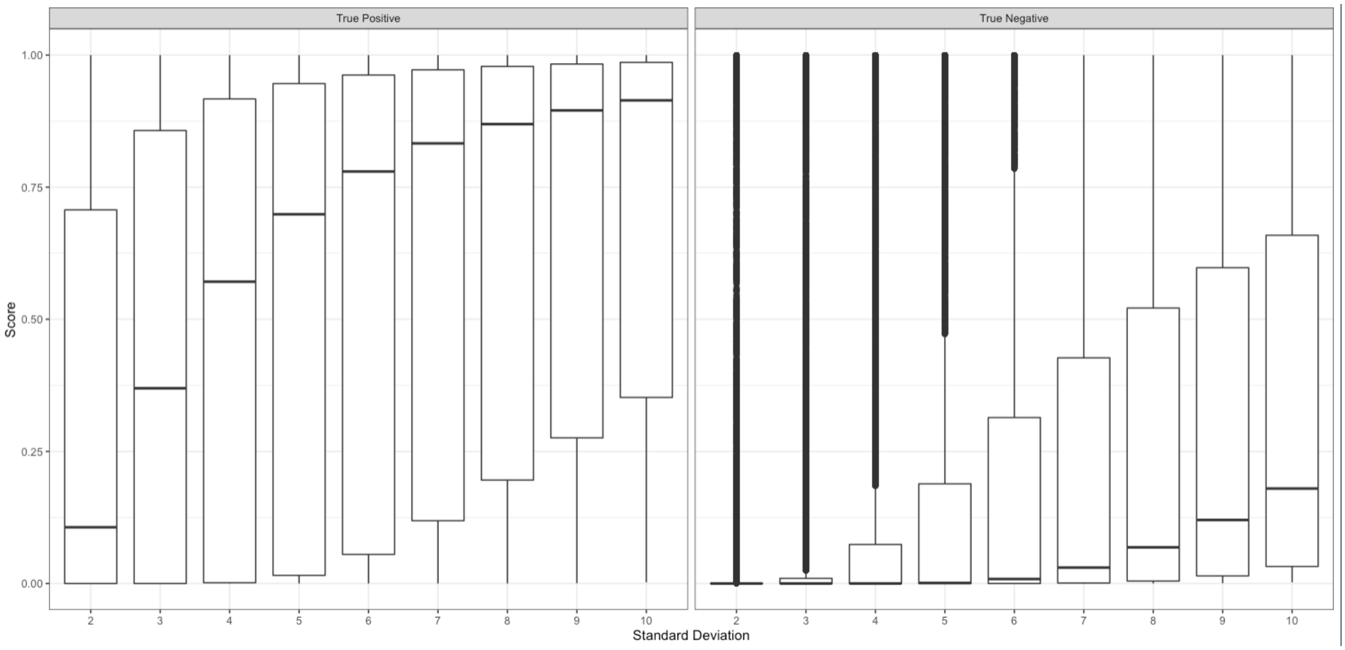


**Figure S2.** Distributions of the retention index score using different standard deviations (retention index search windows) for (left) true positives and (right) true negatives.

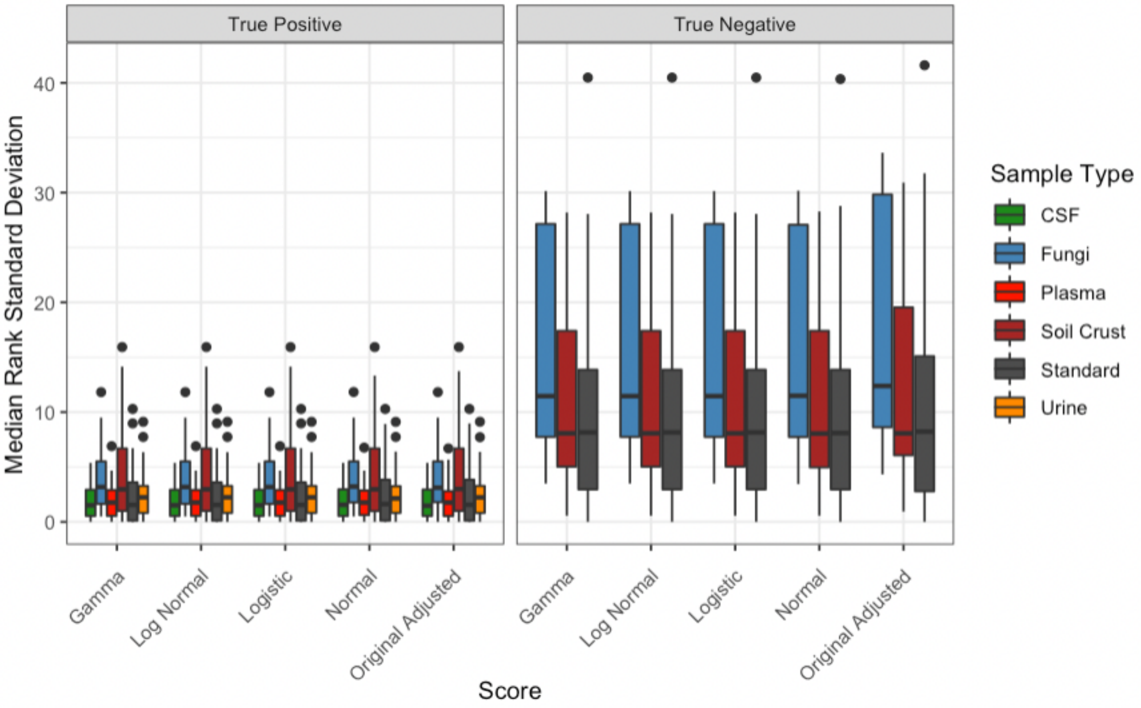


**Figure S3.** The median standard deviation of rank following our 10% holdout analysis that was run 50 times, for all metabolites in our subset (n = 87), broken down by sample type and separated by true positives and negatives.

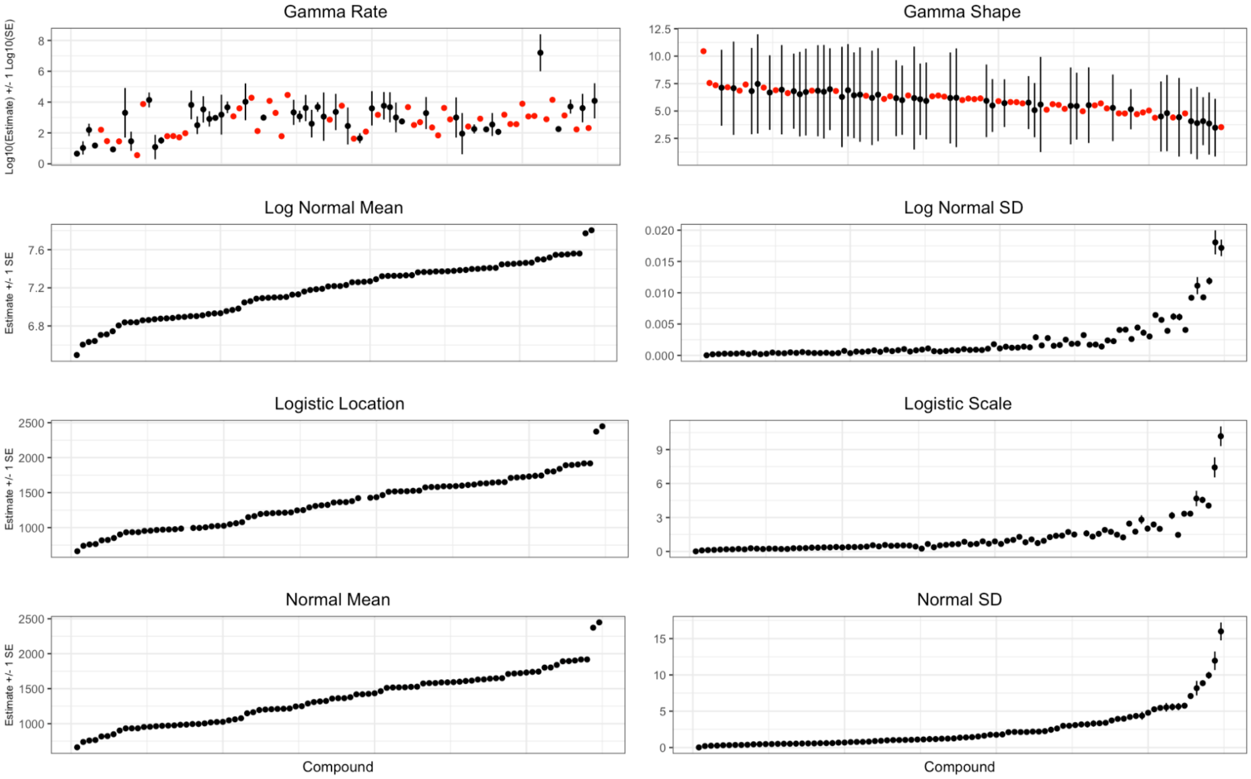


**Figure S4.** The maximum likelihood parameter estimates for each of the four test distributions (top row: gamma, second row: log normal, third row: logistic, last row: normal), where the point indicates the estimation and the lines indicate the standard error, with the exception of gamma which has both of these values log transformed. Red is indicative of a point where the standard error of the estimate failed to calculate.

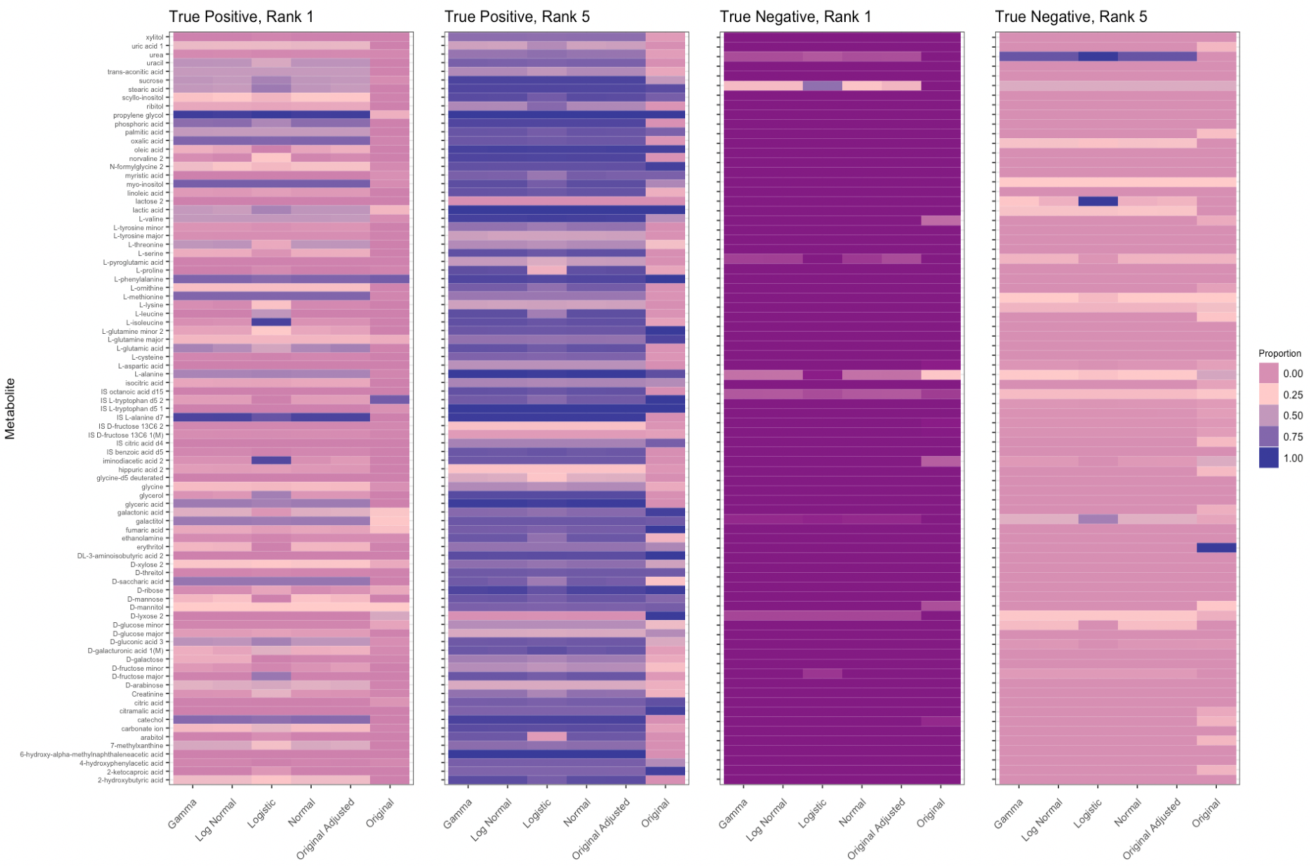


**Figure S5.** Proportions of true positives (left two) and true negatives (right two) at ranks 1 and 5 per metabolite, following the 10% holdout analysis with all input data.

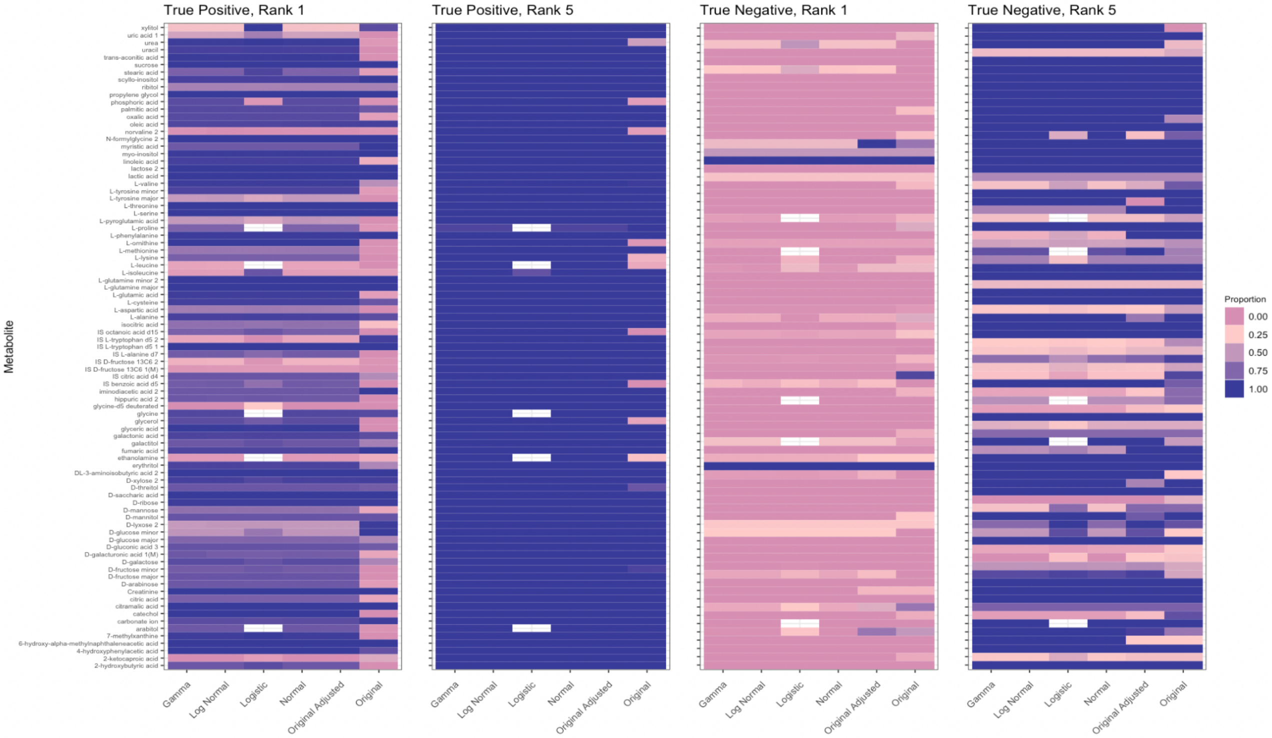


**Figure S6.** Proportions of true positives (left two) and true negatives (right two) at ranks 1 and 5 per metabolite, following the 10% holdout analysis with only metabolites within our subset. A blank value is where the maximum likelihood estimation of the parameters failed in more than 50% of the iterations.

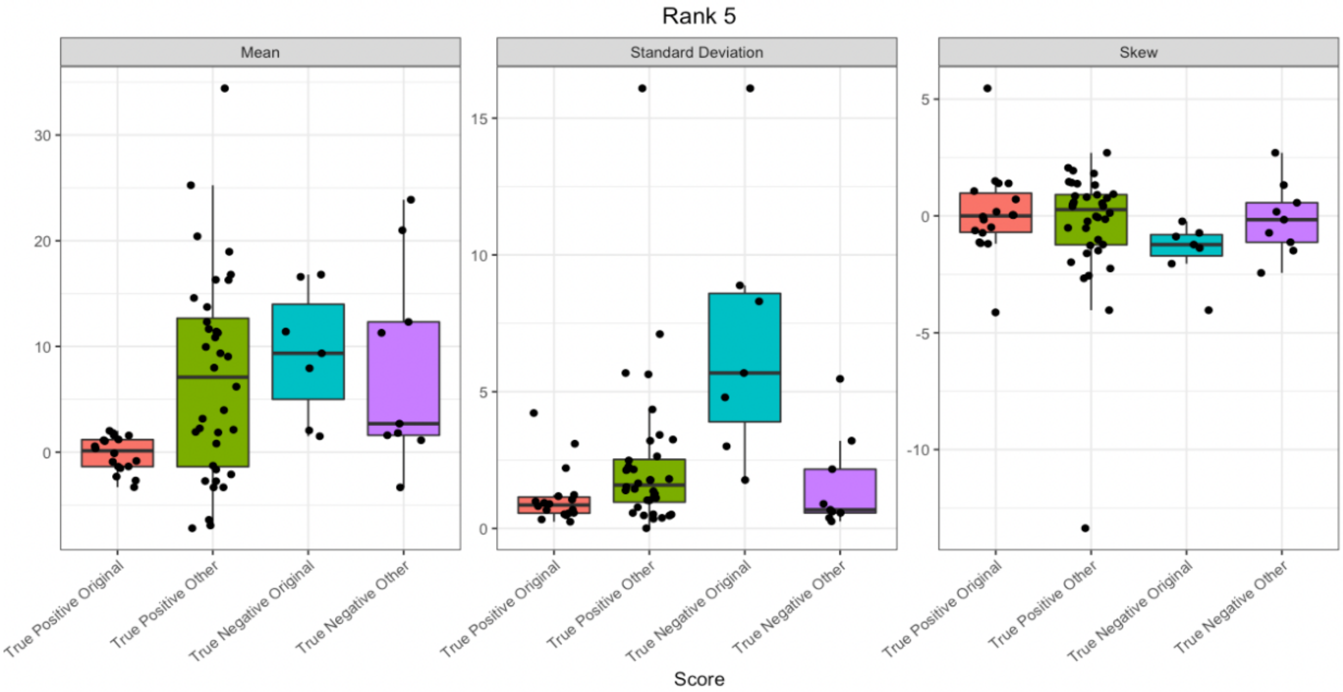


**Figure S7.** The distributional properties of the query retention indices for the best performing scores at rank 5, separated by the metabolites where the original score performed better (true positives: n = 18, true negatives n = 7), or any other score performed better (true positives: n = 36, true negatives n = 9) for the mean and standard deviation. Ties were removed.

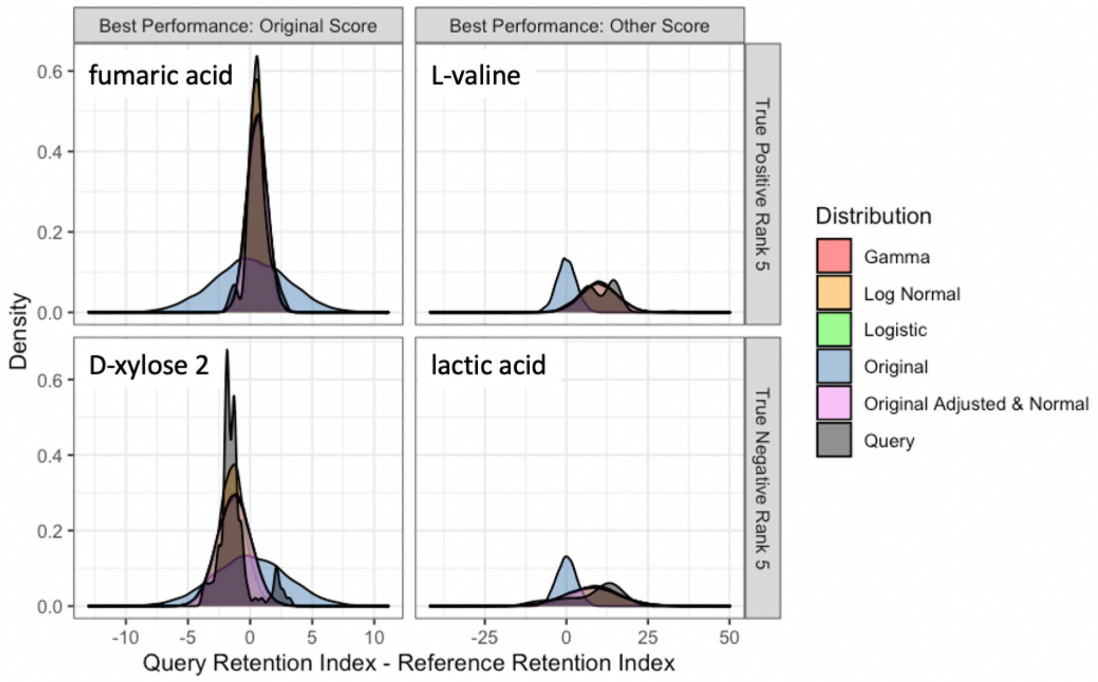


**Figure S8.** Estimated distributions for each score overlaid on the retention index query distribution, centered to each metabolite's reference retention index. Representative metabolites with the best performance (highest true positive proportion or lowest true negative proportion) for the original score (left column) or another distribution (right column) were selected for both true positives (top row) and true negatives (bottom row).
